# Mass Spectrometry-Based Multiomic Profiling Defines Proteome, Lipidome, and Metabolome Remodeling in IFN-γ and LPS-Stimulated BV-2 Microglial Cells

**DOI:** 10.64898/2026.07.27.741024

**Authors:** Abraham M. Borst, Matthew R. Eskritt, Kaitlyn T. Mang, Melissa R. Pergande

## Abstract

Microglial inflammatory activation is accompanied by extensive molecular remodeling, yet proteomic, lipidomic, and metabolomic responses are often analyzed independently. Here, we applied an integrated mass spectrometry-based multiomic workflow to characterize proteomic, lipidomic, and polar metabolomic remodeling from matched BV-2 biological samples following stimulation with interferon-γ and lipopolysaccharide (IFN-γ and LPS). Inflammatory activation was confirmed by increased nitrite accumulation, elevated TNF-α and IL-6 secretion, and treatment-associated morphological changes. Discovery proteomics quantified 8,676 proteins and identified 562 significantly altered proteins, including 344 increased and 218 decreased proteins. Increased proteins were enriched for interferon-responsive, innate immune, inflammatory effector, and antigen-associated pathways, whereas decreased proteins were associated with cellular organization, protein biogenesis, vesicular trafficking, and metabolic regulation. Targeted lipidomics identified 237 significantly altered lipid features out of 356 measured lipids, including increased triacylglycerols and diacylglycerols and broad remodeling of glycerophospholipids, lysophospholipids, and sphingolipid-related species. Targeted polar metabolomics identified 75 significantly altered metabolites out of 98 measured metabolites, including changes in nucleotide/NAD-related metabolism, amino acid metabolism, methylation-associated metabolites, acylcarnitine abundance, phospholipid precursors, polyamine metabolism, arginine/nitric oxide-associated metabolism, and redox-associated metabolites. Process-level integration of significant features revealed coordinated remodeling of inflammatory protein programs with lipid storage, membrane remodeling, nucleotide metabolism, amino acid availability, phospholipid precursor abundance, nitric oxide-associated metabolism, and redox/osmolyte pathways. These findings demonstrate that IFN-γ and LPS-induced activation of BV-2 cells involves integrated immune, lipid, and metabolic adaptation rather than isolated induction of canonical inflammatory mediators. This integrated multiomic framework provides a resource for investigating how lipid and metabolic remodeling regulate microglial inflammatory states.

**Graphical Abstract:** 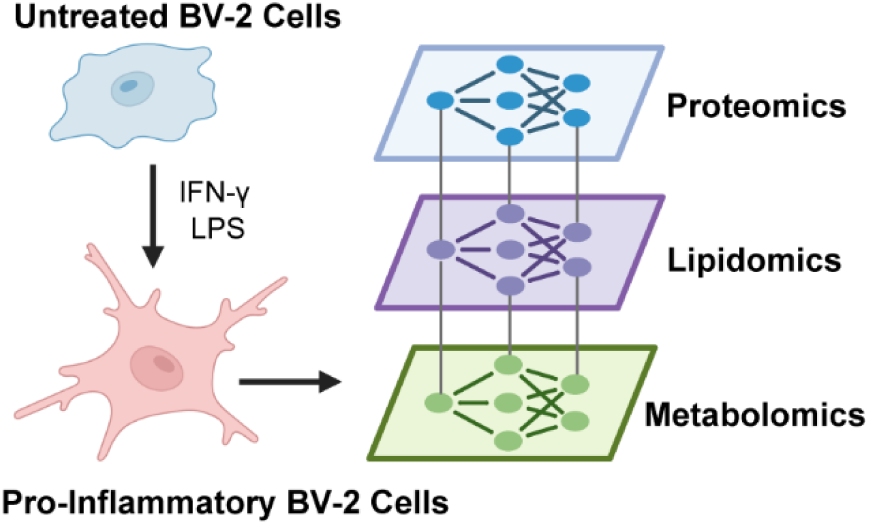

## Introduction

Microglia are the resident immune cells of the central nervous system and play essential roles in immune surveillance, synaptic remodeling, injury response, and maintenance of neural homeostasis.^1^ In response to inflammatory stimuli, microglia undergo rapid functional and molecular remodeling that includes cytokine release, nitric oxide production, altered antigen presentation, metabolic reprogramming, and changes in membrane and lipid signaling pathways.^2–4^ While these responses are critical for host defense and tissue repair, sustained or dysregulated microglial activation is implicated in a broad range of neurological disorders, including neurodegenerative disease, traumatic injury, infection, and neuroinflammatory conditions.^5, 6^ Defining the molecular programs that accompany microglial activation is therefore important for understanding inflammatory signaling in the brain and for identifying pathways that may be therapeutically targeted.

The immortalized BV-2 microglial cell line is widely used as an experimental model for studying microglial inflammatory activation.^7^ Stimulation with interferon-γ (IFN-γ), lipopolysaccharide (LPS), or their combination induces a pro-inflammatory phenotype characterized by increased nitric oxide production, inflammatory cytokine secretion, activation of inflammatory signaling pathways, and broad changes in immune and metabolic function.^8^ Because BV-2 cells are experimentally tractable, reproducible, and compatible with biochemical and molecular profiling workflows, they provide a useful platform for evaluating cellular responses to inflammatory stimuli. However, inflammatory activation is not controlled by a single molecular layer. Instead, it reflects coordinated remodeling of the proteome, lipidome, and metabolome.^9^ A more complete understanding of the BV-2 inflammatory state therefore requires integrated molecular profiling across these complementary layers.

Previous proteomic studies have defined protein abundance changes in activated BV-2 cells, and lipidomic studies have begun to reveal inflammatory remodeling of lipid species and lipid-associated signaling pathways.^10–13^ Although these studies have provided important insight into specific components of the BV-2 response to pro-inflammatory stimulation, most have focused on individual molecular classes, making it difficult to connect immune protein induction with lipid remodeling and metabolic reprogramming in the same experimental system. This is an important limitation because inflammatory microglial activation requires coordinated changes in immune effector proteins, membrane composition, lipid mediators, redox state, amino acid availability, nucleotide metabolism, and energy-associated pathways.^14^ Mass spectrometry-based multiomic workflows are well suited to address this gap because they enable complementary measurement of proteins, lipids, and metabolites from matched biological samples.^15, 16^ Additionally, data-independent acquisition proteomics provides reproducible quantification of thousands of proteins, while targeted lipidomics and metabolomics enable sensitive measurement of lipid classes, lipid species, amino acids, nucleotide-related metabolites, acylcarnitines, redox-associated metabolites, and central metabolic intermediates.^17, 18^ When combined with integrated analysis and pathway enrichment, these approaches can reveal coordinated molecular programs that are not apparent from any single ‘omic layer alone.^19^

In this study, we performed an integrated mass spectrometry-based multiomic analysis of BV-2 cells stimulated with IFN-γ and LPS. We first confirmed inflammatory activation using nitrite accumulation, cytokine secretion, and brightfield morphology. We then applied discovery proteomics, targeted lipidomics, and targeted polar metabolomics to untreated BV-2 samples to define treatment-associated remodeling across molecular layers. Proteomic analysis revealed a robust inflammatory and interferon-responsive signature, including induction of innate immune, antigen-associated, and host-defense pathways. Lipidomic analysis identified broad remodeling of neutral lipids, glycerophospholipids, lysophospholipids, and sphingolipid-related species. Polar metabolomics revealed changes in nucleotide/NAD-related metabolism, amino acid abundance, acylcarnitines, methylation-associated metabolites, glycolytic intermediates, phospholipid precursors, arginine/nitric oxide-associated metabolism, and redox-linked metabolites. Together, these data provide an integrated molecular resource for understanding IFN-γ and LPS-induced inflammatory activation in BV-2 cells and demonstrate the value of multiomic profiling for linking immune protein programs with lipid and metabolic remodeling.

## Materials and Methods

### Reagents and Chemicals

BV-2 murine microglial cells were purchased from Cytion (Heidelberg, Germany). Recombinant mouse interferon gamma (IFN-γ) and lipopolysaccharide (LPS) and were purchased from Thermo Fisher Scientific (Waltham, MA, USA). Fetal bovine serum (FBS), RPMI-1640 medium, and antibiotics were purchased from Fisher Scientific (Hampton, NH, USA). FBS was heat-inactivated at 56 °C before use. Lipidomic internal standards, including standards for acylcarnitines, glycerophospholipids, lysophospholipids, sphingolipids, neutral lipids, cholesterol, and free fatty acids, were purchased from Avanti Polar Lipids (Alabaster, AL, USA), Cayman Chemical (Ann Arbor, MI, USA), Cambridge Isotope Laboratories (Tewksbury, MA, USA), Larodan AB (Solna, Sweden), and Matreya LLC (State College, PA, USA). Stable-isotope-labeled amino acid standards for polar metabolomics were purchased from Cambridge Isotope Laboratories. Unless otherwise specified, all other reagents were purchased from Sigma-Aldrich (St. Louis, MO, USA) and used as received.

### Cell Culture and Microglia Activation

BV-2 cells were maintained in RPMI-1640 medium supplemented with 10% heat-inactivated FBS and 1% penicillin/streptomycin at 37°C in a humidified incubator containing 5% CO₂. To induce a pro-inflammatory microglial phenotype, cells were co-stimulated with IFN-γ and LPS. BV-2 cells were treated with 20 ng/mL IFN-γ and 1 µg/mL LPS for 24 h based on previously reported inflammatory activation conditions for BV-2 microglia.^10, 20^ For proteomic, lipidomic, and metabolomic analyses, BV-2 cells were seeded in 100 mm tissue culture dishes and grown to approximately 75% confluency before treatment. Following 24 h treatment with vehicle or IFN-γ and LPS, BV-2 cells were imaged by brightfield microscopy prior to cell collection to assess treatment-associated changes in cell morphology. Representative fields were acquired using an EVOS M3000 with a 20x objective under identical imaging conditions across both groups. Following imaging, cells were harvested as pellets, flash-frozen on dry ice, and stored at −80 °C until processing. Culture supernatants were collected after 24 h, clarified by centrifugation at 13,000 × g for 10 min, and analyzed immediately for nitric oxide production or flash-frozen and stored at −80 °C for subsequent IL-6 and TNF-α measurements.

### Griess Assay for Nitric Oxide Production

Nitric oxide production was assessed by measuring nitrite accumulation in culture supernatants using a Griess nitrite determination kit (Invitrogen, USA). Clarified culture supernatants were analyzed immediately after collection according to the manufacturer’s protocol. Briefly, 150 µL of sample or nitrite standard was combined with 130 µL of Milli-Q water and 20 µL of Griess reagent in a 96-well plate. The Griess reagent was prepared immediately before use by mixing component A, 0.1% N-(1-naphthyl)ethylenediamine dihydrochloride, and component B, 1% sulfanilic acid in 5% phosphoric acid, at a 1:1 ratio. Plates were incubated at room temperature for 30 min, and absorbance was measured at 548 nm. Nitrite concentrations were calculated from a standard curve ranging from 0 to 100 µM sodium nitrite.

### ELISA Measurement of IL-6 and TNF-α

IL-6 and TNF-α were quantified in clarified BV-2 culture supernatants using mouse-specific ELISA kits from R&D Systems (USA). Assays were performed according to the manufacturer’s instructions. Optical density was measured at 450 nm, and cytokine concentrations were calculated using standard curves generated for each analyte.

### Proteomic Sample Preparation and Mass Spectrometry Analysis

Control and IFN-γ and LPS stimulated BV-2 cell pellets (n=5 per condition) were lysed in 100 µL of 5% SDS supplemented with 1× Halt protease inhibitor (Thermo Fisher Scientific, Waltham, MA, USA). Protein concentrations were determined using a bicinchoninic acid assay, and 20 µg of protein from each sample was used for proteomic sample preparation. Proteins were reduced with 10mM dithiothreitol for 15 min at 55 °C and alkylated with 30mM iodoacetamide for 20 min at room temperature in the dark. Protein digestion and detergent removal were performed using S-Trap Micro spin columns (ProtiFi, Huntington, NY, USA) according to the manufacturer’s protocol; proteins were digested overnight with sequencing-grade trypsin at 37°C using an enzyme-to-protein ratio of 1:40. Peptides were eluted, dried using a CentriVap concentrator (Labconco, Kansas City, MO, USA), and reconstituted in 0.1% formic acid. Peptide concentration was determined using a peptide bicinchoninic acid assay, and 400 ng of peptide from each sample was loaded onto Evotips (Evosep Biosystems, Odense, Denmark).

Peptide samples were analyzed using an Evosep One liquid chromatography system (Evosep Biosystems, Odense, Denmark) coupled to a timsTOF Pro 2 mass spectrometer (Bruker Daltonics, Bremen, Germany). Peptides were separated using the Evosep 30 samples per day method on an EV1137 Performance Column packed with ReproSil-Pur C18, 1.5 µm particles, 15 cm × 150 µm inner diameter (Evosep Biosystems). Mobile phase A consisted of 0.1% formic acid in water and mobile phase B consisted of 0.1% formic acid in acetonitrile. Data were acquired in data-independent mode with parallel accumulation-serial fragmentation (dia-PASEF) over an *m/z* range of 100-1700 and an ion mobility range of 0.75-1.30 1/K₀.

Raw LC-MS/MS files were processed using DIA-NN (version 2.5.1) with library-free searching enabled. Spectral libraries were generated in silico using the reviewed UniProtKB/Swiss-Prot Mus musculus database. Trypsin/P was specified as the protease, with up to one missed cleavage allowed. Carbamidomethylation of cysteine was set as a fixed modification. Precursor and protein identifications were filtered at a 1% false discovery rate. Additional DIA-NN parameters included an *m/z* precursor range of 300-1800, fragment ion range of 200-1800, precursor charge states of 2-5, peptide lengths of 7-30 amino acids, and robust LC high-precision quantification. Protein groups were filtered to require a minimum of two peptides per protein. Protein abundances were log₂ transformed and normalized using variance-stabilizing normalization. Differential protein abundance was assessed using a linear model with group as the explanatory variable and false discovery rates were calculated using the Benjamini-Hochberg procedure. For proteins not observed in one condition, missing group means were estimated using the prolfqua ‘ContrastsMissing’ approach, in which the missing condition mean is approximated using the average abundance at the 5th percentile of the group in which the protein was not quantified.^21, 22^ These missing-contrast cases were annotated separately in the differential-abundance output. Proteins were considered significantly altered if they met both an FDR threshold of <0.05 and an absolute log₂ difference threshold of ≥0.7. Differential abundance results were visualized by plotting differentially expressed proteins.

### Lipidomic and Metabolomic Sample Preparation and Mass Spectrometry Analysis

BV-2 cell pellets were extracted using a Folch-based liquid-liquid extraction to recover lipids and polar metabolites from the same biological samples.^23, 24^ Briefly, cell pellets corresponding to 40 µg of protein were transferred to 1.5 mL microcentrifuge tubes and spiked with class-specific internal standards before extraction. Internal standards include isotopically labeled and non-endogenous standards for acylcarnitines, glycerophospholipids, lysophospholipids, sphingolipids, neutral lipids, cholesterol, bile acids, and free fatty acids, as listed in Table S1. Samples were extracted with 750 µL 2:1 chloroform:methanol, vortexed, and incubated on ice to promote metabolite and lipid partitioning. Phase separation was induced by addition of 250 µL of water followed by centrifugation at 2,000 × g for 5 min at 4 °C. The lower organic phase was collected for lipidomic analysis, and the upper aqueous phase was collected for polar metabolite analysis. Extracts were dried under vacuum.

For lipidomic analysis, dried organic extracts were reconstituted in 50 µL of methanol and analyzed using an Agilent 1290 Infinity II liquid chromatography system coupled to an Agilent 6495 triple quadrupole mass spectrometer equipped with an Agilent Jet Stream electrospray ionization source. Lipids were separated by reversed-phase chromatography using an Agilent ZORBAX Eclipse Plus C18 column (100 × 2.1 mm, 1.8 µm) with mobile phase A consisting of 10 mM ammonium formate, 5 μM Agilent deactivator additive (p/n 5191-3940) in 5:3:2 water:acetonitrile:2-propanol and mobile phase B consisting of 10 mM ammonium formate in 1:9:90 water:acetonitrile:2-propanol. Lipid species were detected using dynamic multiple reaction monitoring in positive and negative ion modes. For polar metabolomic analysis, dried aqueous extracts were reconstituted in 50 µL of 75% acetonitrile, 20% water and 10% methanol, clarified by centrifugation, and analyzed using hydrophilic interaction liquid chromatography coupled to tandem mass spectrometry on the Agilent 6495 triple quadrupole platform. Polar metabolites were separated on an InfinityLab Poroshell 120 HILIC-Z column (2.1 x 150 mm, 2.7 μm) using mobile phase A consisting of 20 mM ammonium acetate containing 5 μM Agilent deactivator and mobile phase B consisting of acetonitrile. Metabolites were detected using scheduled or dynamic MRM transitions in positive and/or negative ion mode. Peak integration was performed using Agilent MassHunter Quantitative Analysis software. Differential lipid and metabolite abundance analysis was performed in R using integrated peak areas. Data were log₂ transformed and compared between IFN-γ and LPS stimulated and control cohorts. Analytes with p < 0.05 were considered significantly altered.

### Functional Enrichment and Integrative Pathway Analysis

Gene Ontology (GO) Biological Process enrichment analysis was performed to identify biological processes associated with significantly altered proteins in IFN-γ + LPS-stimulated BV-2 cells relative to untreated controls. Significantly increased and decreased proteins were analyzed separately using *Mus musculus* as the reference organism. Enriched GO Biological Process terms were retained using a significance threshold of p ≤ 0.05, as determined by the enrichment tool output. This analysis was used to identify biological processes associated with the upregulated and downregulated protein sets.

### Data processing, statistical analysis, visualization and multiomic integration

Proteomic, lipidomic, and polar metabolomic datasets were processed using R-based workflows. All analyses and figure generation were performed in R version 4.5.2. R packages used for data import, processing, statistical analysis, visualization, heatmap generation, and Excel export included readxl, openxlsx, dplyr, tidyr, stringr, ggplot2, ggrepel, pheatmap, tibble, svglite, cowplot, and scales. For proteomics, normalized protein abundance tables generated from DIA-NN output were analyzed in R to compare IFN-γ and LPS-stimulated BV-2 cells with untreated controls. Protein abundances were log₂ transformed and variance-stabilizing normalization was applied prior to statistical analysis. Differential protein abundance was assessed using linear modeling, and p-values were adjusted for multiple comparisons using the Benjamini-Hochberg false discovery rate correction. Proteins were considered significantly altered using an FDR threshold of <0.05 and an absolute log₂ foldchange threshold of >0.7. Principal component analysis, volcano plots, protein count summaries, and heatmaps of differentially abundant proteins were generated from the processed proteomics dataset.

Targeted lipidomics and polar metabolomics datasets were analyzed separately in R. Peak area tables were imported, annotated by treatment group, log₂-transformed, and summarized across biological replicates. Principal component analysis was performed to visualize treatment-associated separation between untreated and IFN-γ and LPS-stimulated samples. Differential abundance analysis was performed for each lipid or metabolite feature, and features with a nominal p-value <0.05 were considered significantly altered. Lipid class summaries were generated by assigning lipid species to their corresponding lipid classes and counting significantly increased and decreased features within each class. For polar metabolomics, significantly altered metabolites were assigned to broad metabolic categories, including nucleotide/NAD-related metabolism, amino acid metabolism, methylation-associated metabolism, acylcarnitine metabolism, phospholipid precursor metabolism, polyamine-associated metabolism, glycolytic intermediates, and redox/osmolyte-associated metabolism. Volcano plots, heatmaps, lipid class summaries and metabolite category summaries were generated from these analyses.

For multiomic integration, significantly altered proteins, lipid features, and polar metabolites were standardized into a common feature-level table containing feature name, ‘omics layer, log₂ fold change, p-value, adjusted p-value/FDR, direction of change, and assigned biological category. Proteins were considered significantly altered using an FDR threshold of <0.05 and an absolute log₂ fold-change threshold of >0.7. Lipid and polar metabolite features were considered significantly altered using a nominal p-value threshold of <0.05. Integrated biological categories were assigned based on feature annotation and known functional relevance. Categories included interferon/host-defense, inflammatory effector, antigen processing/presentation, mitochondrial metabolism, vesicular trafficking/endolysosomal, cytoskeletal organization, ribosome/protein biogenesis, proteostasis/stress response, neutral lipid storage/remodeling, glycerophospholipid remodeling, lysophospholipid remodeling, sphingolipid remodeling, nucleotide/NAD-related metabolism, amino acid metabolism, arginine/nitric oxide-related metabolism, methylation/one-carbon metabolism, phospholipid precursors, acylcarnitine metabolism, polyamine metabolism, glycolytic/TCA intermediates, and redox/osmolyte metabolism.

## Results

### IFN-γ and LPS stimulation induces a pro-inflammatory phenotype in BV-2 cells

BV-2 cells were stimulated with IFN-γ and LPS for to induce a pro-inflammatory state (**Figure 1A**). Brightfield microscopy revealed treatment-associated morphological changes, including an increased proportion of elongated or polarized cells relative to untreated cells (**Figure 1B**). IFN-γ and LPS stimulation also markedly increased nitrite accumulation in the culture medium, consistent with enhanced nitric oxide production (**Figure 1C**), and increased secretion of the pro-inflammatory cytokines IL-6 and TNF-α (**Figure 1D,E**). Together, these results confirmed that IFN-γ and LPS stimulation induced a robust pro-inflammatory phenotype in BV-2 cells suitable for downstream multiomic analysis.

**Figure 1.**
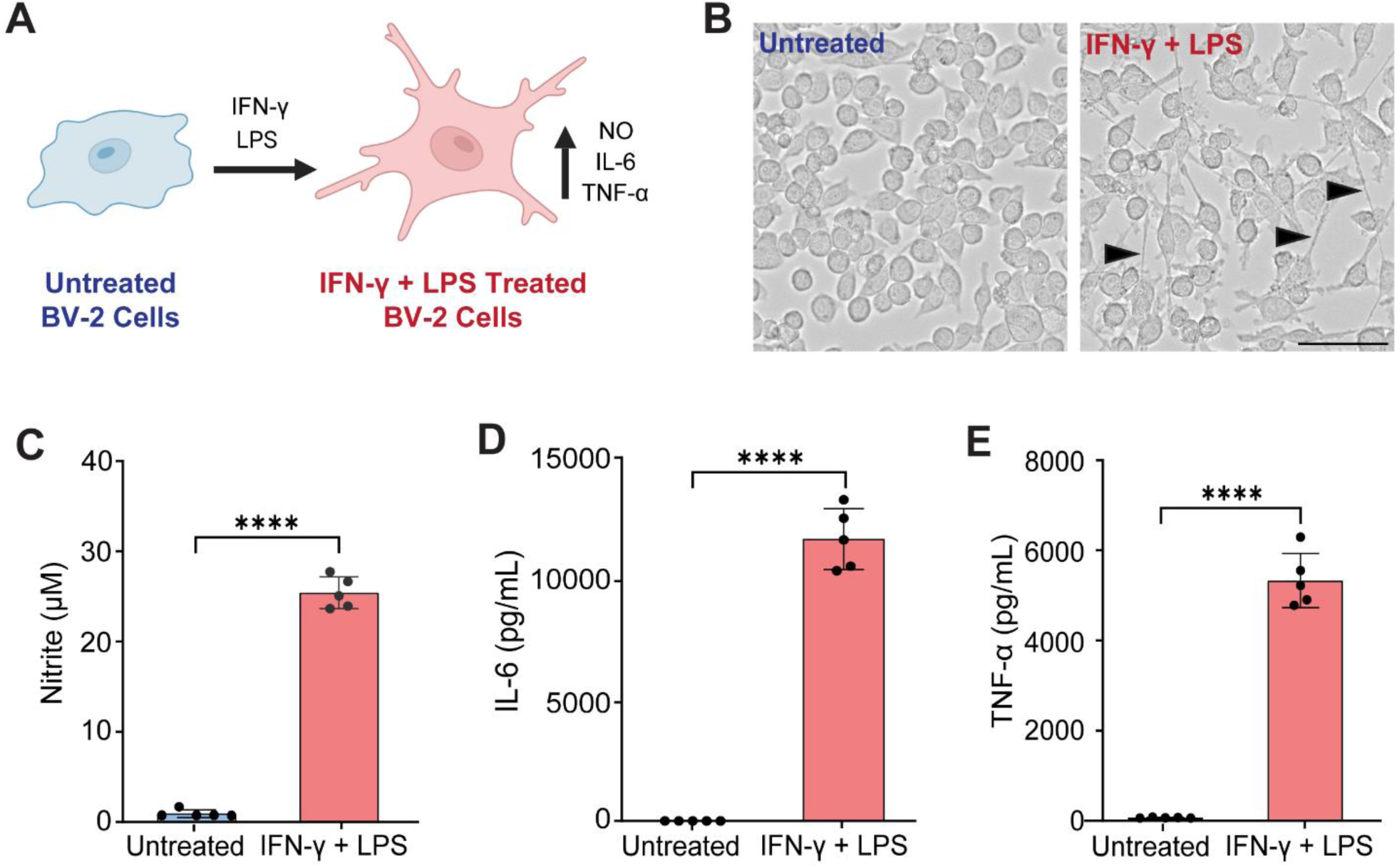
IFN-γ and LPS stimulation induces a pro-inflammatory phenotype in BV-2 microglial cells. (A) Schematic of the experimental model. BV-2 cells were stimulated with IFN-γ and LPS for 24 h to induce a pro-inflammatory phenotype characterized by increased production of nitric oxide and secretion of the inflammatory cytokines IL-6 and TNF-α. (B) Representative brightfield images of untreated and IFN-γ and LPS-stimulated BV-2 cells. Arrowheads indicate elongated or polarized cells observed following stimulation. Scale bar, 50 µm. (C) Nitrite accumulation in the culture medium, used as an indirect measure of nitric oxide production. Concentrations of secreted (D) IL-6 and (E) TNF-α cytokines in the culture medium. Statistical significance was determined using an unpaired two-tailed t-test. ****p < 0.0001.

### Inflammatory activation produces extensive proteomic remodeling in BV-2 cells

Proteomic analysis was performed to characterize protein-level remodeling associated with inflammatory activation. A total of 8,676 proteins were quantified, with similar numbers of protein groups detected across untreated and IFN-γ and LPS-stimulated samples (**Figure 2A**, Table S2). Principal component analysis showed clear separation between the two experimental groups, indicating that inflammatory stimulation produced a reproducible and distinct proteomic profile (**Figure 2B**). Differential abundance analysis identified substantial bidirectional remodeling of the BV-2 proteome following IFN-γ and LPS treatment (**Figure 2C**). Using an FDR threshold of <0.05 and an absolute log₂ fold-change threshold of >0.7, 562 proteins were significantly altered, including 344 increased and 218 decreased proteins. Heatmap visualization of these proteins showed coordinated abundance patterns across biological replicates and clear separation of untreated and stimulated samples (Figure S2). Among the most strongly increased proteins were CMPK2, ACOD1/IRG1, IRGM1, IFI211, and IL1RN, whereas SORL1, VWA8, LTV1, AVIL, and ALDH1L2 were among the prominently decreased proteins. The identities, directions of change, annotated subcellular localizations, and functional relevance of the proteins labeled in the volcano plot are summarized in **Table 1**.

**Figure 2.**
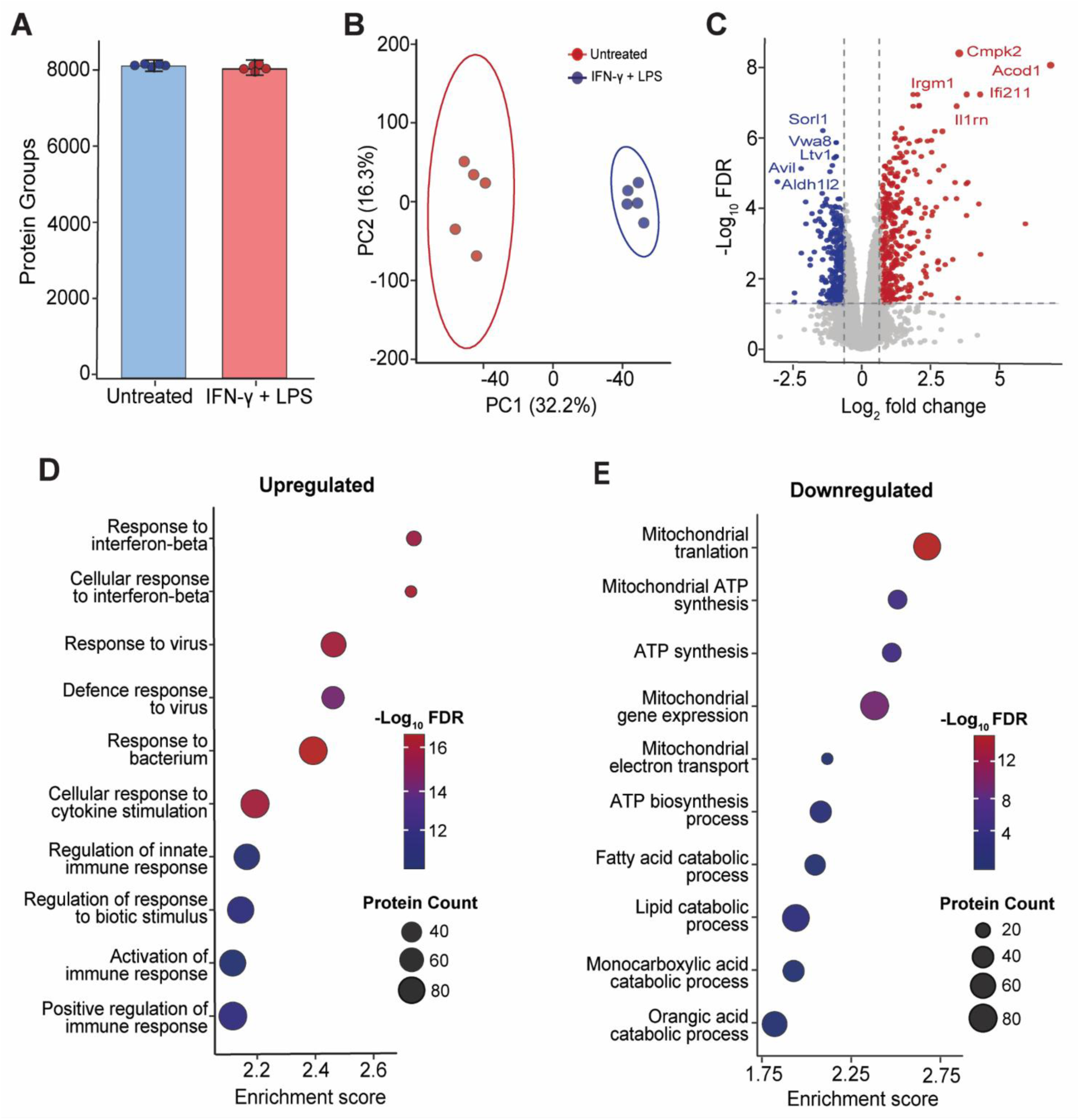
Proteomic remodeling in IFN-γ and LPS-stimulated BV-2 cells. Mass spectrometry-based proteomics was used to quantify global protein abundance changes in untreated and IFN-γ and LPS-stimulated BV-2 cells. (A) Bar graph showing comparable protein groups quantified per condition. (B) Principal component analysis of the proteomics dataset showing separation of untreated and IFN-γ and LPS-stimulated samples. (C) Volcano plot showing differentially abundant protein groups in IFN-γ and LPS-stimulated cells relative to untreated controls. Significantly increased and decreased proteins were defined using an adjusted p-value (FDR) threshold of <0.05 and an absolute log₂ fold change threshold of >0.7. Representative significantly altered proteins are labeled. Gene Ontology biological process enrichment analysis of significantly (D) upregulated and (E) downregulated proteins. Dot size represents the number of proteins associated with each term, and color indicates the adjusted p-value.

**Table 1.**
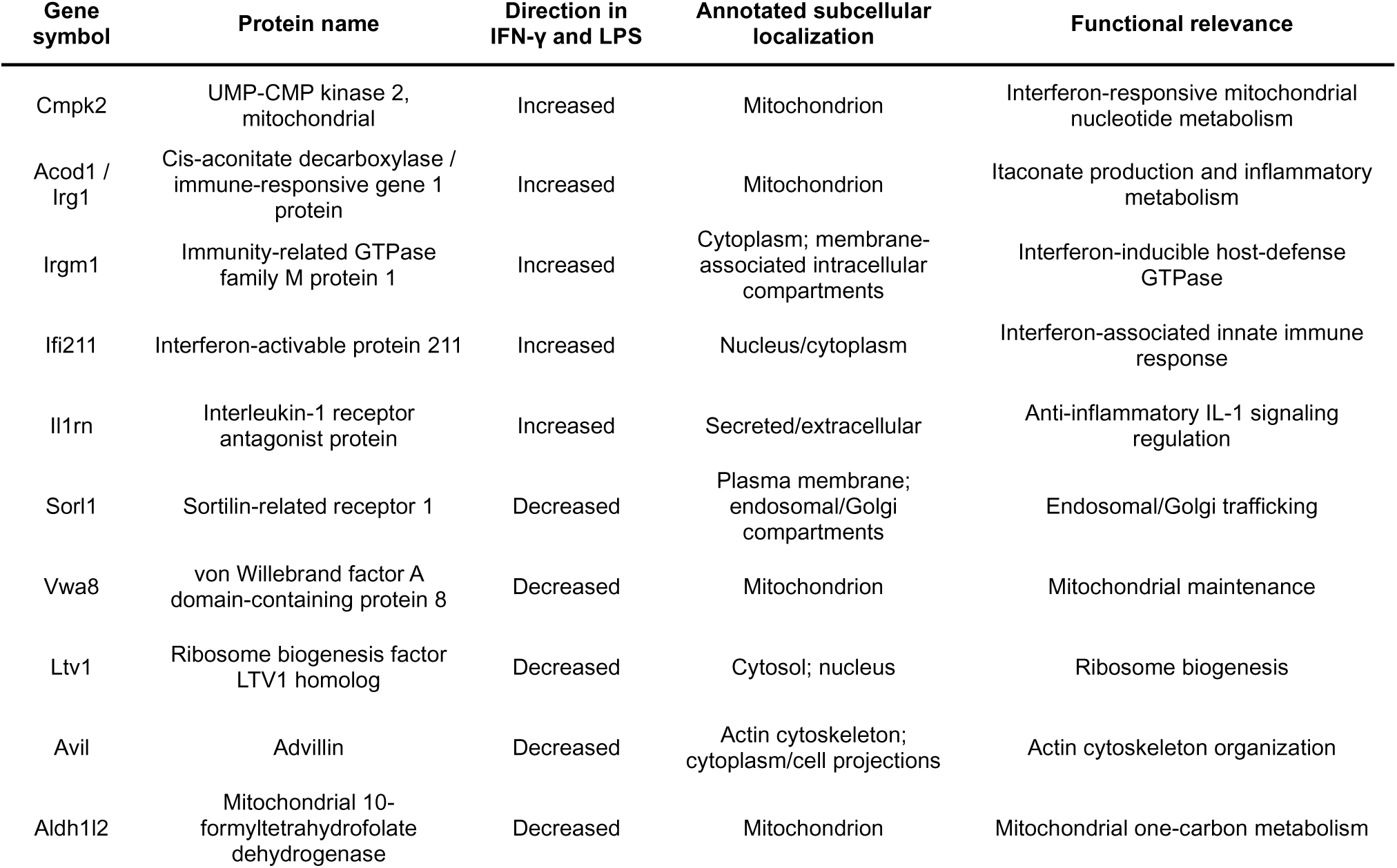
Differentially abundant immune, metabolic, and structural proteins highlighted in IFN-γ and LPS-stimulated BV-2 cells. Proteins listed correspond to the selected significantly altered proteins labeled in the Figure 2C volcano plot comparing IFN-γ and LPS-stimulated BV-2 cells with untreated controls. Direction indicates the abundance change following IFN-γ and LPS stimulation. Annotated subcellular localizations and functional descriptions were assigned from curated mouse protein/gene annotations and published literature.

| Gene symbol | Protein name | Direction in IFN-γ and LPS | Annotated subcellular localization | Functional relevance |
| --- | --- | --- | --- | --- |
| Cmpk2 | UMP-CMP kinase 2, mitochondrial | Increased | Mitochondrion | Interferon-responsive mitochondrial nucleotide metabolism |
| Acod1 / Irg1 | Cis-aconitate decarboxylase / immune-responsive gene 1 protein | Increased | Mitochondrion | Itaconate production and inflammatory metabolism |
| Irgm1 | Immunity-related GTPase family M protein 1 | Increased | Cytoplasm; membrane-associated intracellular compartments | Interferon-inducible host-defense GTPase |
| Ifi211 | Interferon-activable protein 211 | Increased | Nucleus/cytoplasm | Interferon-associated innate immune response |
| Il1rn | Interleukin-1 receptor antagonist protein | Increased | Secreted/extracellular | Anti-inflammatory IL-1 signaling regulation |
| Sorl1 | Sortilin-related receptor 1 | Decreased | Plasma membrane; endosomal/Golgi compartments | Endosomal/Golgi trafficking |
| Vwa8 | von Willebrand factor A domain-containing protein 8 | Decreased | Mitochondrion | Mitochondrial maintenance |
| Ltv1 | Ribosome biogenesis factor LTV1 homolog | Decreased | Cytosol; nucleus | Ribosome biogenesis |
| Avil | Advillin | Decreased | Actin cytoskeleton; cytoplasm/cell projections | Actin cytoskeleton organization |
| Aldh1l2 | Mitochondrial 10-formyltetrahydrofolate dehydrogenase | Decreased | Mitochondrion | Mitochondrial one-carbon metabolism |

The proteins highlighted in **Figure 2C** and **Table 1** illustrate the breadth of the inflammatory proteomic response. Increased abundance of CMPK2, IRGM1, and IFI211 is consistent with activation of interferon-responsive and innate immune pathways.^25–27^ Increased ACOD1/IRG1 further indicates engagement of itaconate-associated immunometabolic remodeling,^28, 29^ while increased IL1RN may represent a compensatory response that limits excessive IL-1 signaling.^30, 31^ In contrast, decreased abundance of SORL1, LTV1, and AVIL suggests remodeling of vesicular trafficking, ribosome biogenesis, and cytoskeletal organization, respectively.^32–34^ Reduced VWA8 and ALDH1L2 further indicate that inflammatory activation is accompanied by changes in mitochondrial maintenance and one-carbon metabolism.^35, 36^ Together, these proteins provide representative examples of immune, metabolic, trafficking, translational, and structural programs remodeled across the BV-2 proteome.

Gene Ontology Biological Process enrichment analysis extended these individual protein observations by showing that the differentially abundant proteins organized into coordinated biological programs. Proteins increased following stimulation were enriched in innate immune activation, interferon-responsive signaling, defense responses, and regulation of inflammatory and cytokine-mediated pathways (**Figure 2D**), consistent with the induction of interferon-inducible proteins and inflammatory metabolic regulators observed in the volcano plot. In contrast, proteins decreased following stimulation were associated with cellular organization, protein synthesis and processing, intracellular trafficking, and metabolic regulation (**Figure 2E**). These downregulated pathways suggest that IFN-γ and LPS activation involves not only induction of immune and host-defense machinery, but also reorganization of constitutive cellular functions. Collectively, the individual-protein and pathway-level changes demonstrate that IFN-γ and LPS stimulation drives a coordinated proteomic transition in BV-2 cells, marked by interferon-driven inflammatory activation together with remodeling of mitochondrial metabolism, protein biogenesis, vesicular trafficking, and cytoskeletal organization.

### Targeted lipidomics reveals extensive remodeling of neutral lipids, phospholipids, and sphingolipids following inflammatory stimulation

Targeted lipidomics was performed to determine whether IFN-γ and LPS stimulation altered lipid pathways associated with membrane composition, inflammatory signaling, or cellular metabolic state (**Figure 3**, Table S3). Principal component analysis of the lipidomics dataset showed separation between IFN-γ and LPS-stimulated and untreated BV-2 cells, with PC1 and PC2 accounting for 60.9% and 18.1% of the variance, respectively (**Figure 3A**). Differential analysis identified 237 significantly altered lipid features out of 356 measured lipids using a nominal p-value threshold of <0.05. Of these, 183 lipid features were increased and 54 were decreased following stimulation (**Figure 3B**). Lipid class-level analysis revealed broad remodeling across multiple lipid families (**Figure 3C**), with the largest numbers of significantly altered lipids observed among phosphatidylethanolamines (PE; 56 altered of 85 measured), triacylglycerols (TG; 34 of 43), phosphatidylcholines (PC; 30 of 42), ceramides (Cer; 21 of 32), sphingomyelins (SM; 18 of 32), phosphatidylinositols (PI; 12 of 14), phosphatidylserines (PS; 11 of 18), diacylglycerols (DG; 9 of 10), and lysophosphatidylcholines (LPC; 9 of 20). Median class-level log₂ fold-change values indicated particularly strong increases in TG, DG, cardiolipin, phosphatidylglycerol, PI, and PS classes, whereas lysophosphatidylethanolamines and hexosylceramides showed an overall decrease.

**Figure 3.**
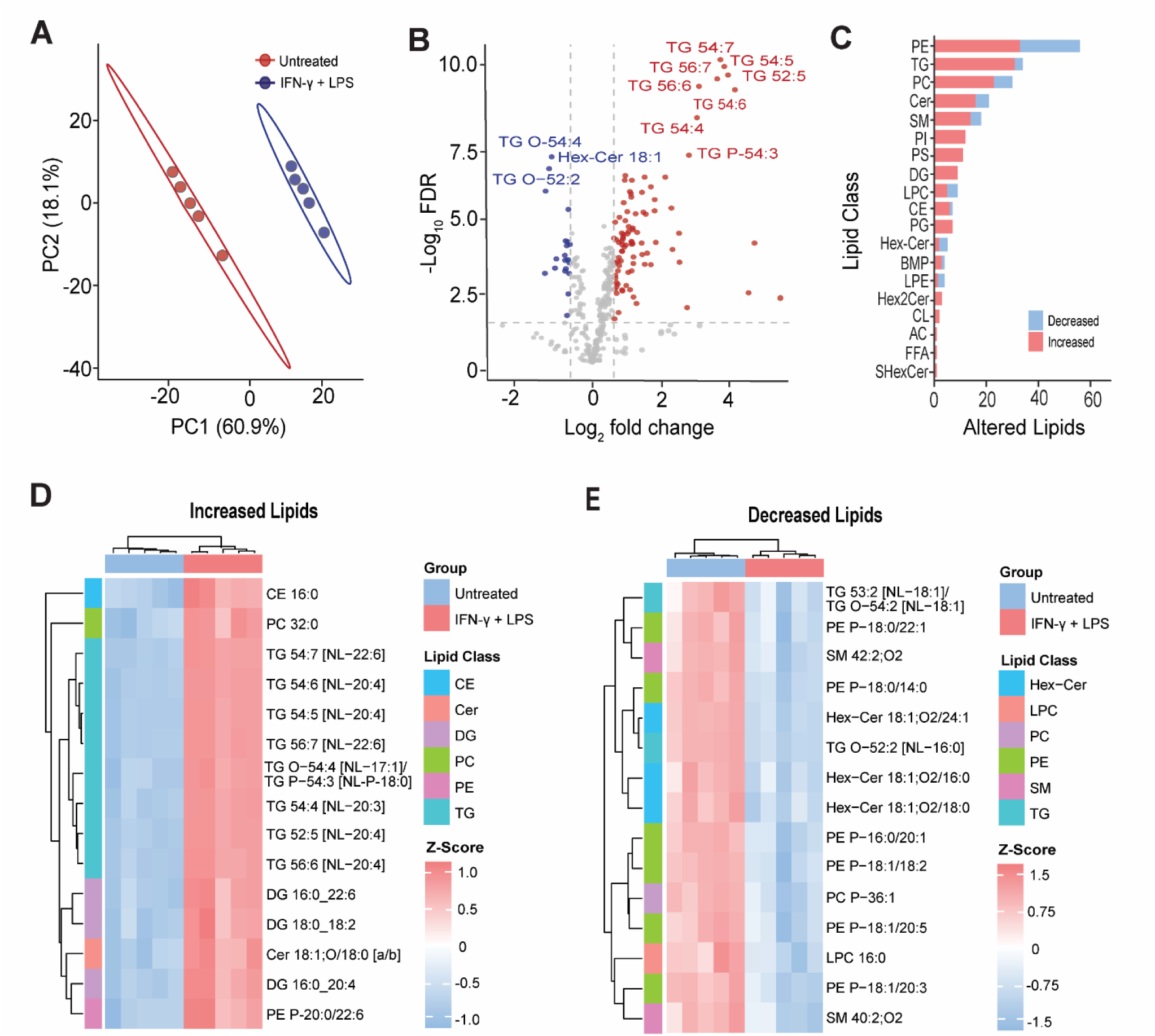
Targeted lipidomics reveals broad lipid remodeling following IFN-γ and LPS stimulation. Targeted lipidomics was used to quantify lipid remodeling in untreated and IFN-γ and LPS-stimulated BV-2 cells. (A) Principal component analysis of the lipidomics dataset showing separation of untreated and IFN-γ and LPS-stimulated samples. (B) Volcano plot of lipid features altered by IFN-γ and LPS stimulation relative to untreated cells. Significantly altered lipids were defined using a nominal p-value threshold of <0.05. (C) Lipid class-level summary showing the number of significantly increased and decreased lipid species within each lipid class. Heatmaps of top-ranked (D) increased and (E) decreased lipid species following IFN-γ and LPS stimulation. Heatmaps show row-scaled log₂-transformed lipid abundance values across biological replicates. The color scale represents the row z-score, where red indicates relatively higher abundance and blue indicate relatively lower abundance within each lipid species.

At the individual lipid-species level, several of the most significantly increased features were polyunsaturated TG species, including TG 54:7, TG 54:5, TG 52:5, TG 56:7, TG 56:6, and TG 54:6 (**Figure 3D**). Several of these TG species showed large increases, with log₂ fold-change values greater than 3, consistent with accumulation or remodeling of neutral lipid pools during inflammatory activation.^37, 38^ Additional increased lipids included DG 16:0_20:4, DG 16:0_22:6, CE 16:0, PC 32:0, and Cer 18:1;O/18:0. In contrast, decreased lipid species included Hex-Cer 18:1;O2/24:1, TG O-52:2, LPE 16:0, LPE 18:1, LPC 16:0, LPC 18:1, SM 42:2;O2, and several plasmalogen PE and PC species (**Figure 3E**). Together, these data indicate that inflammatory stimulation of BV-2 cells produces coordinated remodeling of neutral lipid storage species, glycerophospholipids, lysophospholipids, and sphingolipid-related pathways.

### Polar metabolomics identifies nucleotide, amino acid, acylcarnitine, and redox-associated metabolic remodeling

Targeted polar metabolomics was performed to define metabolic changes associated with inflammatory activation in BV-2 cells (**Figure 4**, Table S3). Principal component analysis showed clear separation between IFN-γ and LPS-stimulated and untreated BV-2 cells, with PC1 and PC2 accounting for 69.3% and 9.1% of the variance, respectively (**Figure 4A**). Differential analysis identified 75 significantly altered metabolites out of 98 measured metabolites using a nominal p-value threshold of <0.05. Of these, 29 metabolites were increased and 46 were decreased following IFN-γ and LPS stimulation (**Figure 4B**). To organize these metabolite changes into broader biological programs, significantly altered metabolites were assigned to major metabolic categories. Category-level analysis showed that IFN-γ and LPS stimulation affected nucleotide/NAD-related metabolism, amino acid metabolism, methylation/one-carbon metabolism, acylcarnitine metabolism, phospholipid precursor metabolism, polyamine metabolism, glycolytic/TCA intermediates, redox/osmolyte metabolism, and arginine/nitric oxide-related metabolism (**Figure 4C**).

**Figure 4.**
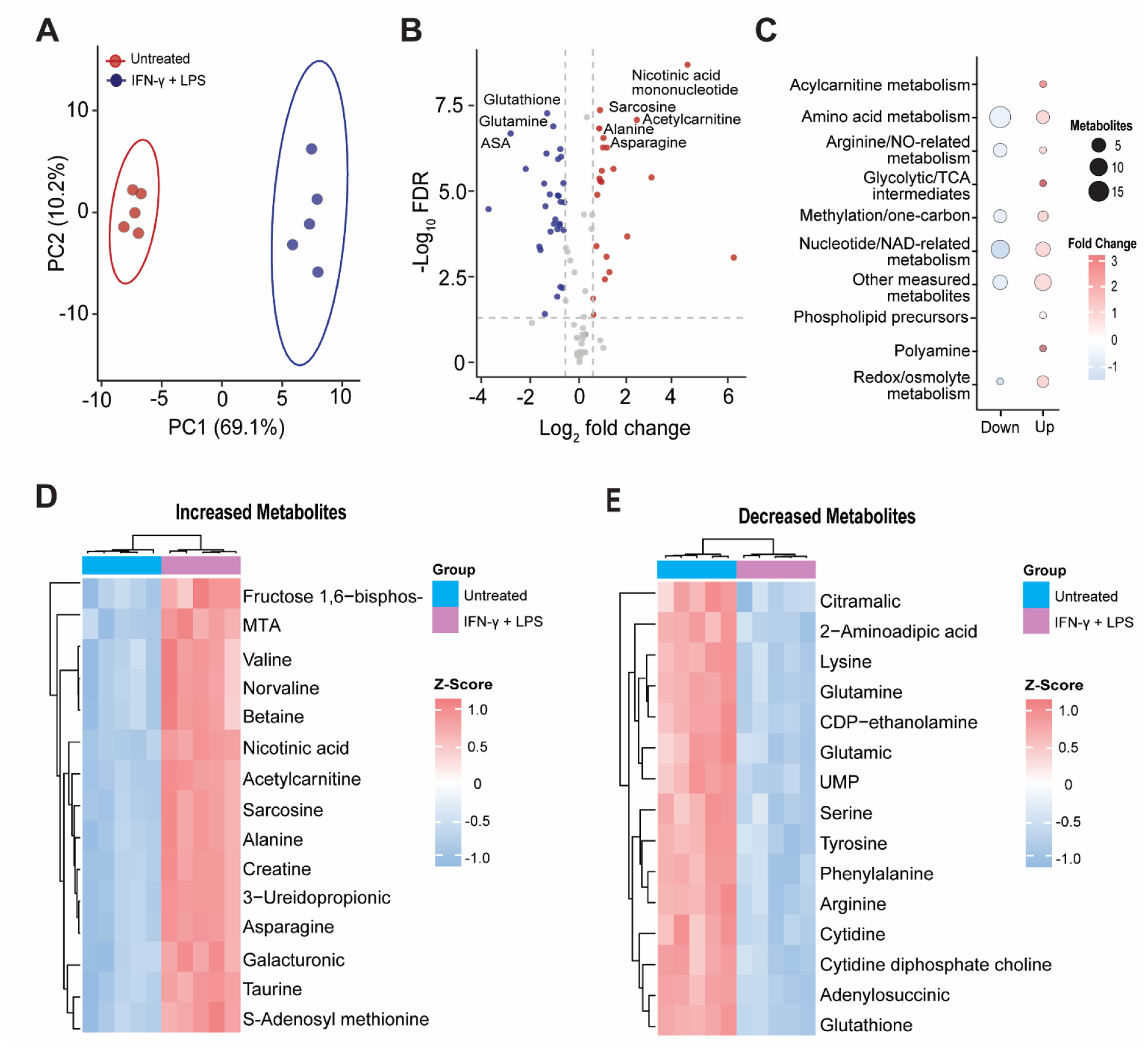
Targeted polar metabolomics identifies metabolic remodeling in IFN-γ and LPS-stimulated BV-2 cells. Targeted polar metabolomics was used to quantify changes in polar metabolites in untreated and IFN-γ and LPS-stimulated BV-2 cells. (A) Principal component analysis of the polar metabolomics dataset showing separation of untreated and IFN-γ and LPS-stimulated samples. (B) Volcano plot shows metabolites altered by IFN-γ and LPS stimulation relative to untreated cells. Significantly increased and decreased metabolites were defined using a nominal p-value threshold of <0.05. (C) Metabolite category bubble plot summarizing significantly altered metabolites by category and direction of change. Bubble size represents the number of significantly altered metabolites in each category, and color indicates the median log₂ fold change. Heatmaps of representative (D) increased and (E) decreased metabolites following IFN-γ and LPS stimulation. Heatmaps show row-scaled log₂ transformed metabolite abundance values across biological replicates. The color scale represents the row z-score, where red indicates relatively higher abundance and blue indicates relatively lower abundance within each metabolite. ASA, adenylosuccinic acid; UMP, uridine monophosphate; MTA, 5-deoxy-5-(methylthio)adenosine.

Heatmap visualization of representative increased and decreased metabolites highlighted coordinated treatment-associated changes across biological replicates (**Figure 4D,E**). Increased metabolites included nicotinic acid mononucleotide, sarcosine, creatine, acetylcarnitine, 3-ureidopropionic acid, asparagine, alanine, S-adenosylmethionine, 5-deoxy-5-(methylthio)adenosine (MAT), taurine, betaine, valine, norvaline, fructose 1,6-bisphosphate, ADP-ribose, and N-acetylputrescine (**Figure 4D**). These changes suggest remodeling of NAD-related metabolism, amino acid metabolism, methylation-associated metabolism, acylcarnitine abundance, glycolytic intermediates, osmolyte/redox-associated metabolites, and polyamine-associated pathways. In contrast, metabolites decreased following IFN-γ and LPS stimulation included oxidized glutathione, glutamine, adenylosuccinic acid, CDP-choline, CDP-ethanolamine, UMP, ADP, inosine, ADMA, 2-aminoadipic acid, lysine, arginine, serine, tryptophan, tyrosine, phenylalanine, cytidine, and UDP-N-acetylglucosamine (**Figure 4E**). These decreases point to broad remodeling of nucleotide metabolism, amino acid availability, phospholipid precursor metabolism, arginine/nitric oxide-related metabolism, and glutathione-associated redox balance. Together, these data demonstrate that IFN-γ and LPS-induced inflammatory activation is accompanied by coordinated remodeling of polar metabolic pathways, including nucleotide/NAD-related metabolism, amino acid metabolism, phospholipid precursor abundance, redox-associated metabolism, and arginine/nitric oxide-associated metabolic changes.^28, 39^

### Multiomic integration reveals coordinated remodeling of inflammatory protein programs, lipid pathways, and metabolic processes

To determine how IFN-γ and LPS stimulation reshaped BV-2 cells across molecular layers, significantly altered proteins, lipids, and polar metabolites were integrated into a shared process-level framework (**Figure 5**, Table S4). This analysis incorporated significant features from the proteomics, lipidomics, and polar metabolomics datasets and assigned them to major biological categories based on molecular annotation and functional relevance. To focus the integrated analysis on interpretable biological programs, features assigned to nonspecific “Other” categories were excluded, and only processes represented by multiple altered molecules were visualized. The resulting integrated bubble heatmap summarizes both the number of altered molecules and the median log₂ fold-change value for each process within each omics layer.

**Figure 5.**
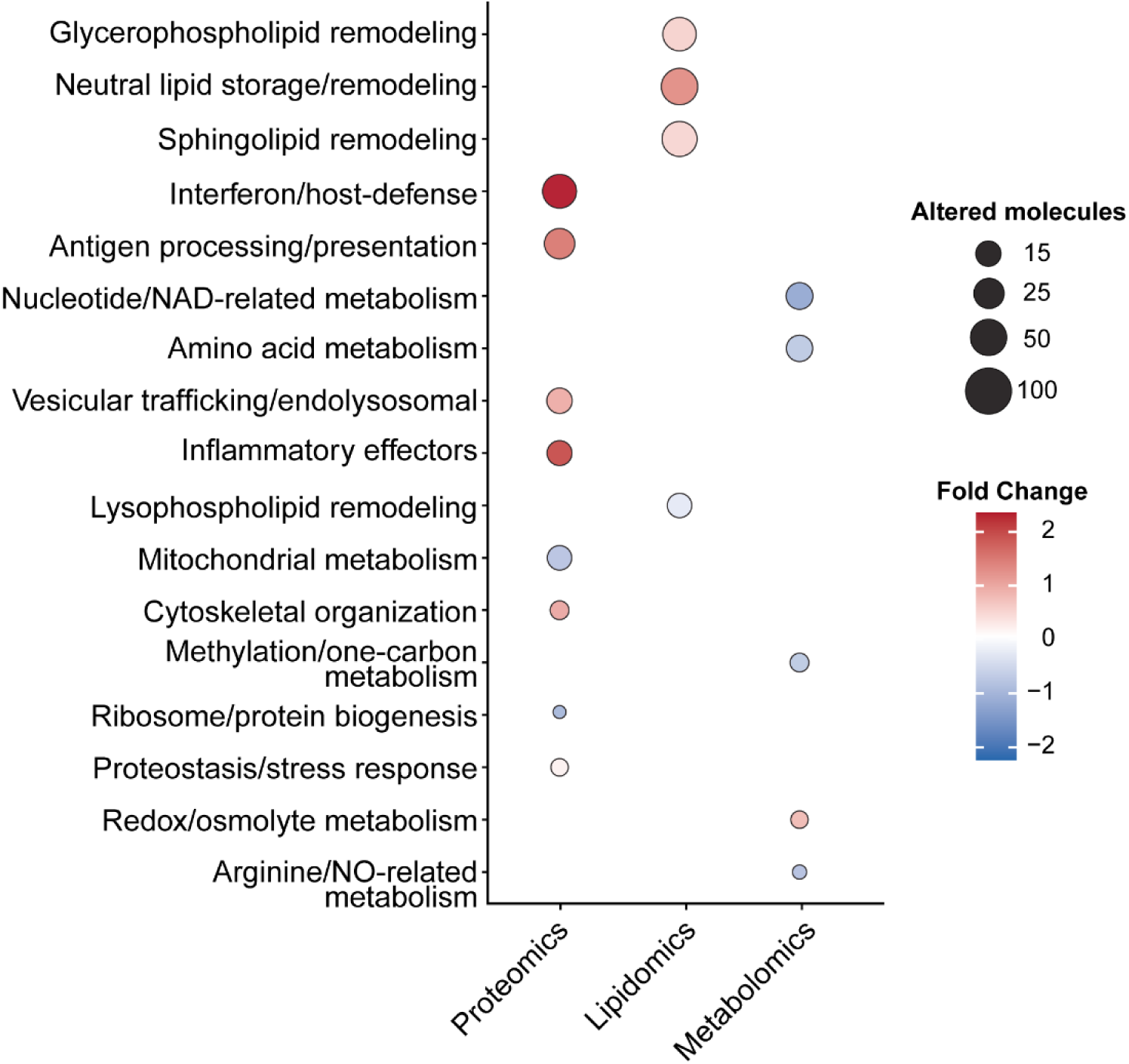
Integrated multiomics analysis reveals coordinated immune, lipid, and metabolic adaptation in IFN-γ and LPS-stimulated BV-2 cells. Significantly altered proteins, lipids, and polar metabolites were integrated to identify biological processes remodeled across the proteome, lipidome, and polar metabolome following IFN-γ and LPS stimulation. Bubble plot summarizing altered molecules by integrated biological process and ‘omic layer. Bubble size represents the number of significantly altered molecules assigned to each biological process within each ‘omic layer. Bubble color represents the median log₂ fold change. Proteins were considered significantly altered using an FDR threshold of <0.05 and an absolute log₂ fold-change threshold of >0.7. Lipid and polar metabolite features were considered significantly altered using a nominal p-value threshold of <0.05.

This process-level integration highlighted coordinated remodeling of immune, lipid, and metabolic programs following inflammatory stimulation. Proteomic changes were most strongly represented by interferon/host-defense proteins, inflammatory effector proteins, and antigen processing/presentation pathways, consistent with established inflammatory and interferon-associated responses in activated microglia.^43–45^ Proteomic changes also included mitochondrial/metabolic proteins, vesicular trafficking/endolysosomal proteins, cytoskeletal organization, ribosome/protein biogenesis, and proteostasis/stress-response pathways. Lipidomic changes mapped primarily to neutral lipid storage/remodeling, glycerophospholipid remodeling, lysophospholipid remodeling, and sphingolipid remodeling. Polar metabolite changes were distributed across nucleotide/NAD-related metabolism, amino acid metabolism, arginine/nitric oxide-related metabolism, methylation/one-carbon metabolism, phospholipid precursor metabolism, acylcarnitine metabolism, polyamine metabolism, glycolytic/TCA intermediates, and redox/osmolyte metabolism.

Together, these cross-layer patterns suggest that IFN-γ and LPS stimulation produced an integrated inflammatory phenotype rather than independent changes within each ‘omic layer. Increased inflammatory and host-defense proteins occurred alongside broad remodeling of membrane-associated and storage lipid pathways, suggesting that immune effector activation was coupled to lipid reorganization. In parallel, changes in mitochondrial/metabolic proteins, proteostasis/stress-response pathways, and ribosome/protein biogenesis were consistent with metabolite-level remodeling of nucleotide/NAD-related metabolism, amino acid availability, acylcarnitine metabolism, redox/osmolyte balance, and phospholipid precursor abundance. The decrease in arginine-related metabolites, together with increased nitrite accumulation and inflammatory protein programs, further supports engagement of nitric oxide-associated metabolism during inflammatory activation. Overall, these results support a model in which BV-2 inflammatory activation involves coordinated immune, lipid, and metabolic adaptation rather than isolated induction of canonical inflammatory mediators.

## Discussion

In this study, we used an integrated mass spectrometry-based multiomic strategy to analyze matched BV-2 biological samples and define coordinated proteomic, lipidomic, and polar metabolomic remodeling associated with IFN-γ and LPS-induced inflammatory activation. While prior studies have examined individual molecular responses to inflammatory stimulation in BV-2 cells and related microglial models, these molecular layers are often analyzed independently. Here, we integrated untargeted proteomics, targeted lipidomics, and targeted polar metabolomics in the same inflammatory BV-2 model. Inflammatory activation was confirmed by increased nitrite accumulation, elevated TNF-α and IL-6 secretion, and treatment-associated morphological changes. Across the three platforms, IFN-γ and LPS stimulation produced extensive molecular remodeling, including 562 significantly altered proteins, 237 significantly altered lipid features, and 75 significantly altered polar metabolites. Together, these datasets show that BV-2 inflammatory activation is not restricted to induction of canonical inflammatory mediators, but instead involves coordinated remodeling of immune protein programs, lipid storage and membrane-associated lipid classes, nucleotide and amino acid metabolism, redox-associated metabolites, phospholipid precursors, and mitochondrial/metabolic processes.

The proteomic response was dominated by proteins associated with interferon signaling, innate immune defense, inflammatory effector function, and antigen-associated pathways. Increased abundance of CMPK2, IRGM1, IFI211, STAT1, PTGS2, CYBB, guanylate-binding proteins, and related interferon-associated factors is consistent with induction of a broad inflammatory and host-defense proteome.^25–27, 30, 31^ Increased ACOD1/IRG1 was particularly notable because this enzyme links inflammatory activation to itaconate-associated immunometabolic remodeling, while increased IL1RN may reflect induction of cytokine-regulatory feedback pathways.^28, 29^ In parallel, decreased abundance of proteins linked to vesicular trafficking, ribosome biogenesis, cytoskeletal organization, mitochondrial function, and one-carbon metabolism suggests that IFN-γ and LPS stimulation also reorganizes constitutive cellular programs. These findings are consistent with the Gene Ontology enrichment results, which showed enrichment of immune and interferon-associated processes among increased proteins and enrichment of cellular organization, protein synthesis and processing, intracellular trafficking, and metabolic regulation among decreased proteins.

The lipidomics and polar metabolomics data demonstrated that inflammatory activation was accompanied by extensive remodeling of lipid and metabolic pathways. Targeted lipidomics revealed increased abundance of multiple triacylglycerol and diacylglycerol species, together with broad changes in glycerophospholipids, lysophospholipids, and sphingolipid-related species. These changes suggest remodeling of neutral lipid pools, membrane composition, phospholipid turnover, and lipid signaling pathways during inflammatory activation.^38,40–43^ Targeted polar metabolomics showed coordinated changes in nucleotide/NAD-related metabolism, amino acid abundance, methylation-associated metabolites, acylcarnitine metabolism, glycolytic intermediates, phospholipid precursors, polyamine-associated metabolites, and redox-linked metabolites. Increased nicotinic acid mononucleotide and ADP-ribose suggest remodeling of NAD-related metabolism, while decreased arginine together with increased nitrite accumulation supports engagement of nitric oxide-associated metabolism during inflammatory activation.^44–46^ Together, these lipid and metabolite changes indicate that IFN-γ and LPS stimulation couples inflammatory signaling to broad lipid and metabolic adaptation.

A major strength of this study is that the multiomic design enabled inflammatory activation to be interpreted as a coordinated molecular response across complementary proteomic, lipidomic, and metabolomic layers, rather than as isolated changes within each dataset. Proteomics defined the immune and stress-response programs induced by IFN-γ and LPS, including interferon/host-defense proteins, inflammatory effectors, antigen-associated pathways, and changes in mitochondrial, trafficking, cytoskeletal, and protein biogenesis pathways. Lipidomics revealed how this inflammatory state was accompanied by remodeling of neutral lipid storage species, membrane-associated glycerophospholipids, lysophospholipids, and sphingolipid-related pathways. Polar metabolomics further connected these changes to altered nucleotide/NAD-related metabolism, amino acid abundance, phospholipid precursor availability, arginine/nitric oxide-associated metabolism, and redox/osmolyte balance. By integrating these datasets into a shared process-level framework, we identified coordinated molecular programs that would be less apparent from any single ‘omic layer alone. Increased inflammatory and host-defense proteins occurred alongside lipid and metabolite changes related to membrane remodeling, lipid storage, redox balance, and metabolic adaptation, supporting a model in which BV-2 inflammatory activation involves coordinated immune, lipid, and metabolic remodeling rather than independent changes within each molecular layer. Importantly, this integration is descriptive rather than causal; it identifies cross-layer relationships and candidate pathways for future validation but does not establish direct mechanistic relationships among proteins, lipids, and metabolites.

Several limitations should be considered. BV-2 cells are an immortalized microglial cell line and do not fully recapitulate the biology of primary microglia or human microglia in vivo. Therefore, the molecular changes identified here should be interpreted as a controlled model of inflammatory activation rather than a complete representation of microglial responses in the brain. In addition, the lipidomics and metabolomics analyses were targeted, meaning that unmeasured lipid species and metabolites may also contribute to the inflammatory response. The lipid and polar metabolite analyses used a nominal p-value threshold, which is appropriate for exploratory targeted profiling but should be followed by validation of key metabolites and lipid species in independent experiments. Finally, the multiomic integration used curated process-level categories to summarize significant molecular changes. This approach improves biological interpretability but does not replace pathway-specific mechanistic experiments, quantitative flux measurements, or direct causal testing.

Overall, this study demonstrates that IFN-γ and LPS stimulation drives broad, coordinated proteomic, lipidomic, and metabolomic remodeling in BV-2 cells. The inflammatory phenotype was characterized by increased nitric oxide and cytokine production, induction of interferon-responsive and innate immune proteins, extensive neutral lipid and membrane lipid remodeling, and widespread changes in nucleotide, amino acid, redox, phospholipid precursor, and mitochondrial-associated metabolism. The multiomic framework established here provides a resource for future studies investigating how lipid and metabolic remodeling regulate microglial inflammatory states.

## Conclusions

This study demonstrates that IFN-γ and LPS stimulation drives coordinated proteomic, lipidomic, and metabolomic remodeling in BV-2 cells. Inflammatory activation was marked by increased nitrite production, elevated IL-6 and TNF-α secretion, induction of interferon-responsive and innate immune protein programs, and remodeling of proteins associated with mitochondrial function, vesicular trafficking, protein biogenesis, and cytoskeletal organization. Targeted lipidomics and polar metabolomics further revealed changes in neutral lipid storage species, glycerophospholipids, lysophospholipids, sphingolipid-related pathways, nucleotide/NAD-related metabolism, amino acid abundance, phospholipid precursor metabolism, arginine/nitric oxide-associated metabolism, and redox-linked metabolites. By integrating these datasets into a shared process-level framework, this work shows that inflammatory activation in BV-2 cells reflects coordinated immune, lipid, and metabolic adaptation rather than isolated changes within individual molecular layers. These findings provide a multiomic resource for future studies of microglial immunometabolism, lipid remodeling, and neuroinflammatory signaling.

## Data Availability

All data contained within this manuscript is available upon request of the corresponding author. All mass spectrometry data have been deposited into the MASSive data repository with the dataset identifier MSV0000XXXXX.

## Supporting information

SupportingInformation

## Acknowledgements

This work was supported by the Department of Chemistry at Boston University. Financial support from Undergraduate Research Opportunities Program (Boston University) to A.M.B. and the Mason award (Boston University) to K.T.M. is gratefully acknowledged.

## Supporting Information

Figure S1 Brightfield morphology of control and IFN-γ and LPS stimulated BV-2 cells S-3

Figure S2 Heatmap of differentially expressed proteins S-4

Table S1 Internal standards used for lipidomic and metabolomic analyses S-5

Table S2 Proteomic analysis

Table S3 Targeted lipidomic and metabolomic analysis

Table S4 Multiomics integration analysis

