## SupportingInformation for "Mass Spectrometry-Based Multiomic Profiling Defines Proteome, Lipidome, and Metabolome Remodeling in IFN-γ and LPS-Stimulated BV-2 Microglial Cells"

### Supporting Information


| | | |
|---|---|---|
| Figure S1 | Brightfield morphology of control and IFN-γ and LPS stimulated BV-2 cells | S-3 |
| Figure S2 | Heatmap of differentially expressed proteins | S-4 |
| Table S1 | Internal standards used for lipidomic and metabolomic analyses | S-5 |
| Table S2 | Proteomic analysis | |
| Table S3 | Targeted lipidomic and metabolomic analysis | |
| Table S4 | Multiomics integration analysis | |

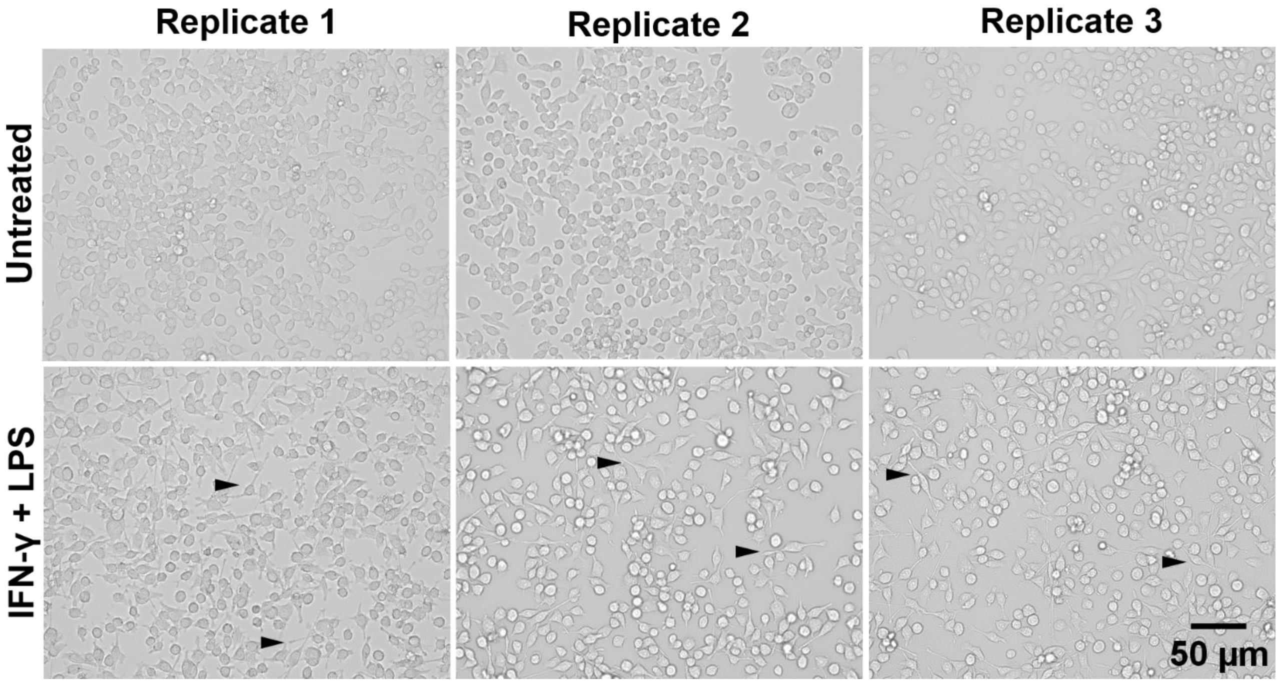


**Figure S1. Brightfield morphology of untreated and IFN-γ and LPS stimulated BV-2 cells.** Representative brightfield images of BV-2 cells following 24 h treatment with vehicle (untreated) or IFN-γ and LPS. Untreated cells displayed typical BV-2 morphology, whereas IFN-γ and LPS cells showed treatment-associated morphological changes, including increased numbers of elongated or polarized cells. Black arrowheads indicate representative elongated morphologies observed in the stimulated condition. Scale bar, 50 µm.

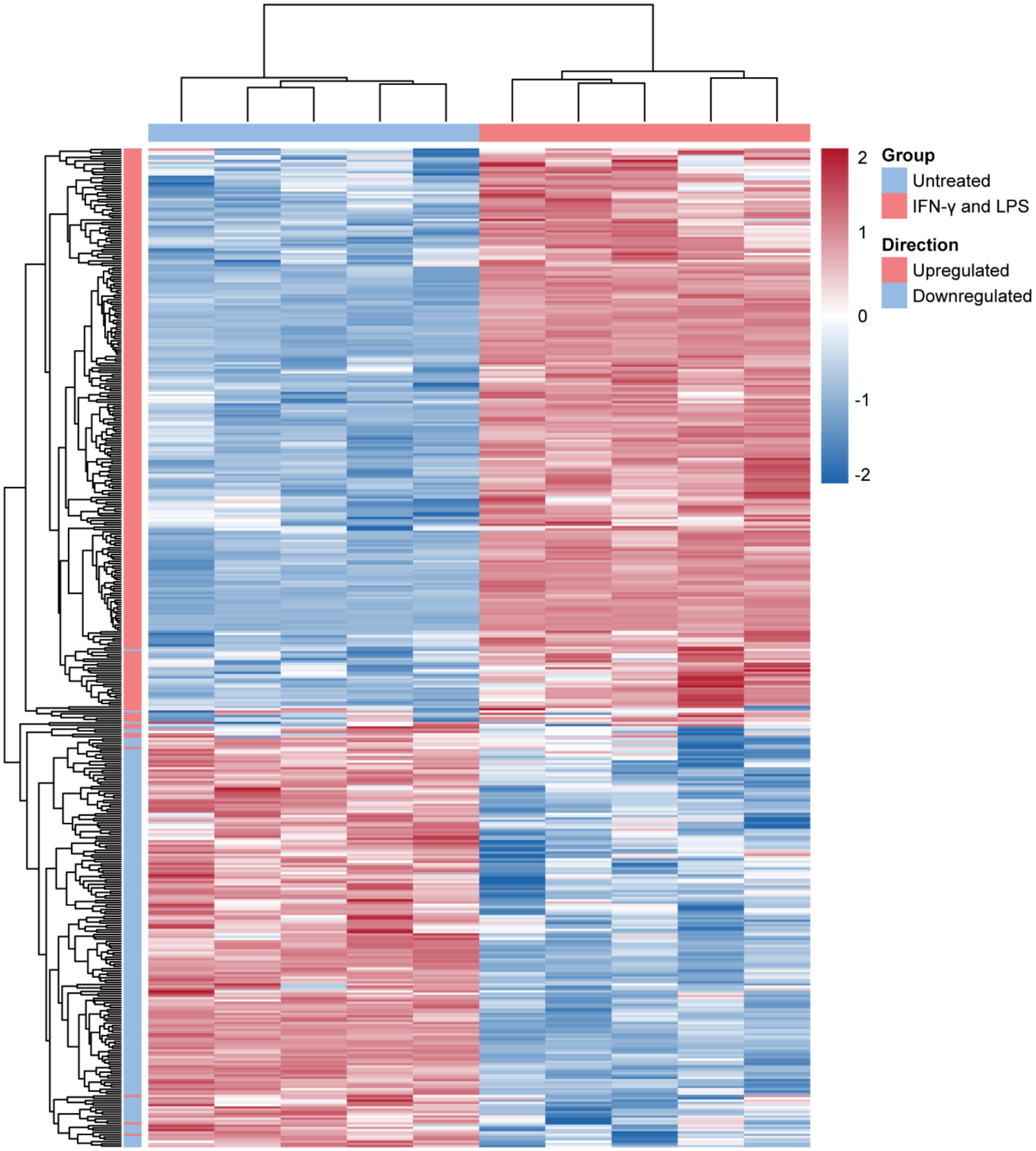


**Figure S2. Heatmap of differentially abundant proteins in IFN-γ and LPS-stimulated BV-2 cells.** Heatmap showing the relative abundance patterns of significantly altered proteins identified by mass-spectrometry-based proteomics in untreated and IFN-γ and LPS-stimulated BV-2 cells. Proteins were considered significantly altered using an adjusted FDR threshold of <0.05 and an absolute $\log_2$ fold-change threshold of >0.7. Rows represent differentially abundant proteins and columns represent biological replicates. Protein abundances are shown as row-scaled normalized intensity values, highlighting treatment-associated increases and decreases across the BV-2 proteome. The heatmap demonstrates coordinated separation of untreated and IFN-γ and LPS-stimulated samples and supports broad proteomic remodeling following inflammatory activation.

| Internal Standard | Class | Vendor | Catalog Number | Stock Concentration (μM) | Volume (μL) per 1 mL Extraction Buffer |
|---|---|---|---|---|---|
| L-Alanine-$d_4$ | Amino Acid | Cambridge Isotope Laboratories | NSK-A-1 | 500 | 1.0 |
| L-Arginine-$^{13}C_1$,$d_4$ · HCl | | | | 500 | |
| L-Aspartic acid-$d_3$ | | | | 500 | |
| L-Citrulline-$d_2$ | | | | 500 | |
| DL-Glutamic acid-$d_3$ | | | | 500 | |
| Glycine-$^{13}C_1$,$^{15}N$ | | | | 2500 | |
| L-Leucine-$d_3$ | | | | 500 | |
| L-Methionine-$d_3$ | | | | 500 | |
| L-Ornithine-$d_6$ · HCl | | | | 500 | |
| L-Phenylalanine-$^{13}C_6$ | | | | 500 | |
| L-Tyrosine-$^{13}C_6$ | | | | 500 | |
| L-Valine-$d_8$ | | | | 500 | |
| AC 16:0 d9 | Acylcarnitine | Avanti Research | 330379 | 24 | 2.0 |
| PC 15:0_18:1 d7 | Glycerophospholipid / lysophospholipid | Avanti Research | 330707 | 212 | 10.0 |
| PE 15:0_18:1 d7 | Glycerophospholipid / lysophospholipid | | | 7 | |
| PS 15:0_18:1 d7 | Glycerophospholipid / lysophospholipid | | | 6 | |
| PG 15:0_18:1 d7 | Glycerophospholipid / lysophospholipid | | | 39 | |
| PI 15:0_18:1 d7 | Glycerophospholipid / lysophospholipid | | | 12 | |
| PA 15:0_18:1 d7 | Glycerophospholipid / lysophospholipid | | | 10 | |
| LPC 18:1 d7 | Glycerophospholipid / lysophospholipid | | | 47 | |
| LPE 18:1 d7 | Glycerophospholipid / lysophospholipid | | | 10 | |
| CE 18:1 d7 | Neutral lipid / sterol | | | 532 | |
| MG 18:1 d7 | Neutral lipid / sterol | | | 6 | |
| DG 15:0_18:1 d7 | Neutral lipid / sterol | | | 17 | |
| TG 15:0_18:1_15:0 d7 | Neutral lipid / sterol | | | 68 | |
| SM 18:1 d9 | Sphingolipid | | | 41 | |
| Cholesterol d7 | Neutral lipid / sterol | | | 254 | |
| w-6 FFA 20:4 d11 | Free fatty acid | Cayman Chemical | 10006758 | 3169 | 1.0 |
| SPBP (S1P) 18:1 d7 | Sphingolipid | Avanti Research | 860659P | 431 | 1.0 |
| Cer 18:0;O2/8:0 | Sphingolipid | Avanti Research | 860626P | 400 | 1.5 |
| SPB 17:1;O2 | Sphingolipid | Avanti Research | 860640P | 400 | 1.0 |

| | | | | | |
|---|---|---|---|---|---|
| Cer d18:1;O2/18:0 d7 | Sphingolipid | Avanti Research | 860677P | 400 | 1.5 |
| Hex2Cer 18:1;O2/15:0 d7 | Sphingolipid | Avanti Research | 330727 | 400 | 1.5 |
| HexCer 18:1;O2/15:0 d7 | Sphingolipid | Avanti Research | 330729 | 400 | 1.5 |
| Hex3Cer 18:1;O2/17:0 | Sphingolipid | Cayman Chemical | 24876 | 400 | 1.5 |
| SHexCer 18:1;O2/12:0 | Sphingolipid | Avanti Research | 860573P | 400 | 1.0 |
| LPI 13:0 | Glycerophospholipid / lysophospholipid | Avanti Research | 850101 | 200 | 1.0 |
| PC P-18:0/18:1 d9 | Ether phospholipid / plasmalogen | Avanti Research | 852475C | 1280 | 1.0 |
| PE P-18:0/18:1 d9 | Ether phospholipid / plasmalogen | Avanti Research | 852474C | 1353 | 1.0 |

**Table S2. Internal standards used for lipidomic and metabolomic analyses.**
